# A flexible anatomic set of mechanical models for the organ of Corti

**DOI:** 10.1101/760835

**Authors:** Jorge Berger, Jacob Rubinstein

## Abstract

We build a flexible platform for the study of the mechanical performance of the organ of Corti (OoC) in the transduction of basilar membrane (BM) vibrations to motion of an inner hair cell bundle (IHB). In this platform, each anatomic component of the OoC is described by an equation of motion that can be followed in time. We propose an initial set of models that attempt to capture the nonlinearities of somatic and bundle motility, but can nevertheless be easily handled. The anatomic components that we consider are the outer hair cells (OHCs), the outer hair cell bundles, Deiters cells, Hensen cells, the IHB and various sections of the reticular lamina. We study endolymph fluid motion in the subtectorial gap and then the mutual interactions among the components of the OoC, including the pressure exerted by endolymph. Minute bending of the apical ends of the OHCs can have a significant impact on the passage of motion from the BM to the IHB, including possible critical oscillator behaviour, even without the assistance of tectorial motion, shearing, or bundle motility. Thus, the components of the OoC could cooperate to enhance frequency selectivity, amplitude compression and signal to noise ratio in the passage from the BM to the IHB. Our models also provide a mechanism that could contribute to appropriate amplification of the wave travelling along the cochlea.

## 1. Introduction

Hearing in mammals involves a long chain of transductions (1–7). Pressure oscillations are collected from the air by the outer ear, and used by the middle ear to shake perilymph in the inner ear, while reducing the impedance mismatch. The wavelength of sound in perilymph is longer than the entire cochlea, but the partitioned structure of the cochlea extracts from it a travelling surface wave with shrinking wavelength, that deposits most of its energy at a short segment of the partition (8, 9). Most of the elastic energy delivered to the cochlear partition resides at the basilar membrane (BM).

We will focus on a slice of the organ of Corti (OoC), that senses the vibrations at a particular position in the BM, transmits them to the corresponding inner hair cell bundle (IHB), and from there to the auditory nerve. From the present point of view, motion of the BM will be the ‘input,’ and motion of the IHB, the ‘output.’ Accordingly, in this treatment the OoC does not include the BM. The shape of the OoC in the basal region of the cochlea is quite different than the shape near the apex; we will have in mind the OoC in the basal region, where higher frequencies are detected, and where the OoC has the greatest impact on amplification and frequency selectivity.

Figure 1 is a schematic drawing (not to scale) of a slice of the OoC, showing the components with which we will deal. It should be noted that whereas the outer hair cell bundles (OHBs) are attached to the tectorial membrane (TM), the IHB is not. As a consequence, when a cuticular plate [the top of an outer hair cell (OHC)] rises, the corresponding OHB tilts clockwise; on the other hand, motion of the reticular lamina (RL) has no direct effect on the inclination of the IHB. In order to turn the IHB and send a signal to the auditory nerve, endolymph flux in the subtectorial channel is required.

**Fig. 1.**
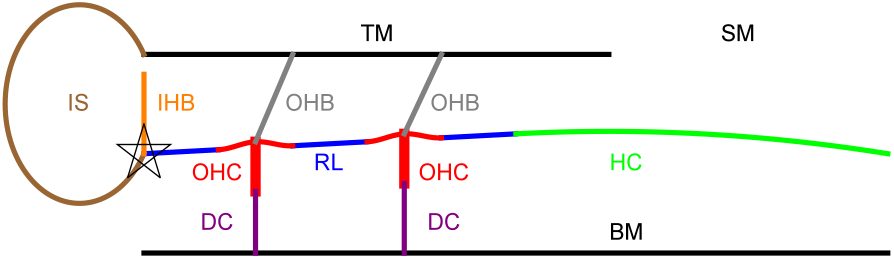
Schematic drawing, showing the components of the OoC. TM: tectorial membrane; SM: scala media; IS: inner sulcus; IHB: inner hair cell bundle; OHB: outer hair cell bundle; OHC: outer hair cell; RL: reticular lamina (set of blue segments); HC: Hensen cells; DC: Deiters cell; BM: basilar membrane. The top of each OHC will be called ‘cuticular plate’ (CP). The model for each of these components is spelled out in Section 3. The star marks the position that is taken as the origin, *x* = *y* = 0.

Our aspiration is not to obtain accurate values for the mechanical performance of the OoC, but rather to gain insight on how its components cooperate to achieve this performance. We would like to answer questions such as: Why the IHB is not attached to the TM? Or, why after transforming fluid flow into mechanical vibration, this vibration is transformed back into fluid flow, this time along a narrow channel, involving high dissipation. Is there any advantage of having several OHCs, rather than a single stronger OHC? How does an OHC perform mechanical work on the system? Is there any role to passive components such as the Hensen cells (HC)?

Many theoretical treatments fall into an extreme category. At one extreme, mechanical activity of the OoC is substituted by an equivalent circuit, and it’s not clear where Newton’s laws come in. At the other extreme, the OoC is divided into thousands of pieces, and a finite elements calculation is carried out (10–14). Neither of these approaches enables us to answer the questions above. Our approach involves postulating a simplified model for each component, with idealised geometry and with as few elements and forces as possible, hoping to capture the features that are essential for its functioning. After the models are chosen, Newton laws can be meticulously followed. Judging from our results, in sections 4 and 5 we conjecture plausible answers to most of the questions above.

Moreover, we would like to be in a position that enables us to investigate broader questions, such as: Could nature have built the OoC, or could an artificial OoC be designed in a different way? In particular, we would like to look for possible mechanisms to achieve frequency tuning and amplitude compression. Substantial evidence has led to the conclusion that the OoC compresses the amplitudes and tunes the frequencies of the vibrations transferred from the stapes to the BM. By taking motion of the BM as the input, we will be mainly investigating the more controversial question of whether there could be an alternative or additional filter that provides compression and tuning on the way from the BM to the auditory nerve (15–20). The conjecture of such a “second filter” is usually attributed to motion of the TM, but our analysis indicates that this feature is not necessary.

Models of the OoC abound (11, 13, 14, 16, 21–23). We do not intend to compete with existing models or to improve them. Rather, we consider complementary aspects. Although our models may be oversimplifications, there are features that are given here more attention than usual. These include the relative motion of different parts of the RL, flow of endolymph along the subtectorial channel and the forces that it exerts.

The most important difference between our models and those we have found in the literature is that pressure in the subtectorial channel is a function of position and time, that exerts large forces along the RL. Another salient difference is that the RL is not regarded as a completely rigid body, but rather the cuticular plates (CPs) can form mild bulges or dents in response to the local forces exerted by the corresponding OHC and OHB.

## 2. Analytical Procedure

### A. Scope and conventions

We deal with a slice of the OoC, so that our analysis is at most two dimensional. Whenever we mention mass, force, moment of inertia, torque, or flow rate, it should be understood as mass (or force, etc.) per unit thickness of the slice. Our set of models is sufficiently simple to permit analytic integrations over space, and we will be left with a system of differential equations for functions of time, that can be solved numerically. Since these equations are nonlinear, we do not perform a Fourier analysis. There are normally three rows of outer hair cells, but we believe that the important fact is that there is more than one, and include just two outer hair cells in our explicit models.

Guided by measurements that indicate that the RL pivots as a rigid beam around the pillar cells head (24, 25), we take the origin at this pivot point. We will assume that the equilibrium positions of the RL and of the upper border of the HC lie along a straight line, that will be taken as the *x*-axis (that will be enviewed as “horizontal” and the *y*-axis will point “upwards”). Traditionally (26) it has been assumed that the relative motion between the TM and the RL, which governs OHC excitation and generates endolymph flow, is predominantly shearing motion. On the other hand, Nowotny and Gummer (27) found that the subtectorial gap can shrink and expand. Recent measurements (in the apical region) (20, 28) obtained that the *x*- and the *y*-component of this relative motion have similar amplitudes. Here we will focus on the pulsatile mode, which is usually disregarded. Accordingly, except for rotational and fluid motion, motion will be restricted to the *y*-direction.

By “height” of the RL, the HC, or the TM, *y*_RL_(*x,t*), *y*_HC_(*x,t*), and *y*_T_(*x,t*), we will imply a position at the surface that is in contact with the endolymph. The width of the subtectorial channel is *D*(*x,t*) = *y*_T_(*x,t*) − *y*_RL_(*x,t*) [or *y*_T_(*x,t*) − *y*_HC_(*x,t*)], and we will assume that in equilibrium *D*(*x,t*) is constant and denote it by *D*_0_. Vertical forces will be considered positive when they act upwards and angular variables will be positive when counterclockwise.

**Fig. 2.**
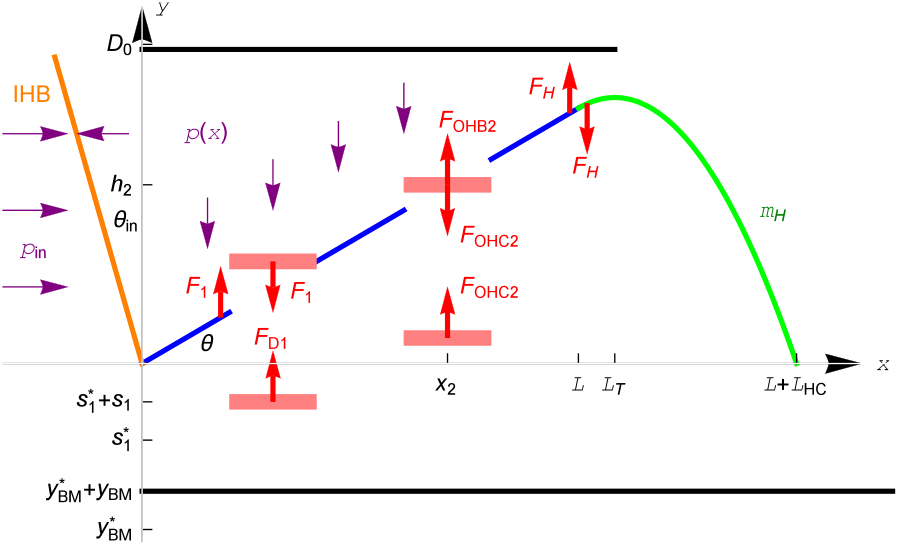
Force diagram (not to scale), showing the movable parts in our model and several of the forces that act on them. Each pink rectangle represents a mass *m*. To avoid clutter, analogous quantities that are present in both OHCs are shown in only one of them. The force between a CP and the RL is 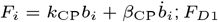 is shorthand for 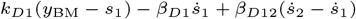; the force exerted by an OHB, *F*_OHA*i*_, is given by Eq. [7]; the tension of an OHC, *F*_OHC*i*_, is given by Eqs. [8] and [9]; the force between the RL and the HC, *F*_H_, can be evaluated using Eq. [11]. 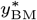 and 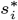 are, respectively, the resting heights of the BM and of an OHC-DC interface, and are not required in our equations.

### B. Common notations and units

We denote by *L, L*_HC_ and *L*_T_ the lengths of the RL, the HC, and the TM. *θ* will be the angle of the RL with respect to the *x*-axis and *θ*_in_ the angle of the IHB with respect to the *y*-axis. We assume that |*θ*(*t*)| ≪ 1, so that the projections of the RL and the HC onto the *x*-axis also cover lengths *L* and *L*_HC_. Several of the coordinates and forces in our models are illustrated in Fig. 2.

For an arbitrary function *f* of position and time, we denote *f*′: = ∂*f*/*∂x* and 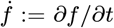. The absolute value of *f*(*t*) at a given time will be denoted as |*f*(*t*)| (with the argument written explicitly), whereas |*f*| will denote the amplitude of *f*, as defined in Appendix 1A.

Pressure exerted by endolymph on OoC components is expected to have a major influence on their motion. Since flow of endolymph is scaled by the height *D*_0_ of the subtectorial gap, it is natural to express all quantities in units that involve *D*_0_. The unit of length will be *D*_0_, the unit of time, 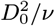, and the unit of mass, 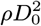, where *ν* and *ρ* are the kinematic viscosity and the density of endolymph. The expected orders of magnitude of these units are *D*_0_ ~ 10*μ*m, 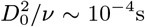, and 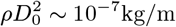. All our variables and parameters will be expressed in terms of these units. Using these units might permit to scale results among cochleae of different sizes.

## 3. Detailed Modelling

We aim at building a flexible platform in which each anatomic component of the OoC is described by a simple model that translates into a simple differential equation. As required, it should be possible to change the model of any of the components, leading to a change in just one of the differential equations in the system. In this way, we can readily check how a given feature in the model affects the performance of the entire OoC. Accordingly, the models below may be regarded as initial guesses. Some of them may capture the behaviour of the component that they represent, and others may not.

A mathematica code that integrates our system of differential equations is available at notebookarchive.org (29). This code is modular, so that not only the parameters can be varied, but also the models.

### A. Subtectorial channel

We denote by *p*(*x,y,t*) the pressure in the endolymph and by *υ*(*x,y,t*) the *x*-component of the local velocity. The flow rate in the *x*-direction is

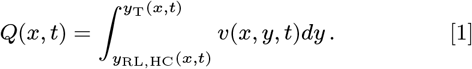

We will assume that motions of the RL, the HC and the TM are very small in comparison to *D*_0_, so that the limits of integration can be set as 0 and *D*_0_ (i.e. 1 in our units). We assume that endolymph is incompressible, so that the net flow entering a region has to be compensated by expansion of that region and therefore

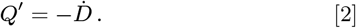

Invoking incompressibility and the fact that the Reynolds number is very small, the *x*-component of the Navier–Stokes momentum equation can be linearised and reduced to

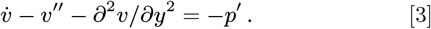

By means of a suitable expansion in powers of *D*_0_/*L* (Appendix 2) we conclude that the pressure can be taken as independent of *y* and obtain the approximate relation

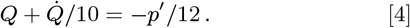

We assume that the only input is motion of the BM, whereas the pressure *p*(*L*_T_) at the exit to the scala media (SM) is taken as constant. We will set *p*(*L*_T_) = 0, i.e., pressure in the SM will be taken equal to the pressure in the tissues under the RL and the HC.

### B. Reticular lamina

We regard it as a straight beam, but exclude the CPs from it, in order to explore the possibility that they bend. The RL obeys the rotational equation of motion

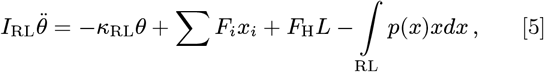

where *I*_RL_ and *κ*_RL_ are the moment of inertia and the rotational stiffness of the RL, *F_i_* is the force exerted on the RL by the CP centered at *x* = *x_i_, F*_H_ is the force exerted on the RL by the HC, and the integration is over the range 0 ≤ *x* ≤ *L* excluding the CPs.

### C. Cuticular plates

CPs are actin rich areas in the apical region of hair cells, where stereocilia bundles are enrooted. In reptiles and amphibians, the cytoplasma between a CP and the surrounding RL has scarce actin filaments and little mechanical resistance (30–32). In mammals, the CP has a lip that protrudes beyond the OHC cross section and extends to adherens junctions with neighboring cells. The *β*-actin density in the CP is much lower than that in stereocilia or in the meshwork through which stereocilia enter the plate, and is therefore expected to be relatively flexible (33). We will assume that each CP can form a bulge (or indentation) relative to the RL. The length of each CP will be *ℓ* and its height *y_i_*(*x*) = *θx* + *b_i_*(1 + cos[2*π*(*x* − *x_i_*)/*ℓ*]), where *b_i_* is the average height above the RL, as illustrated in Fig. 3. Attributing to the CP a mass *m* and a position *y_i_* = *h_i_*: = *θx_i_* + *b_i_*, its equation of motion is

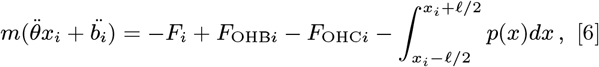

where *F*_OHB*i*_ is the force exerted by the hair cell bundle and *F*_OHC*i*_ the tension of the cell. We set 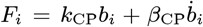, where *k*_CP_ and *β*_CP_ are a restoring and a damping coefficient. The usual assumption that the CPs are fixed within the RL amounts to taking infinite values for *k*_CP_ and *β*_CP_.

**Fig. 3.**
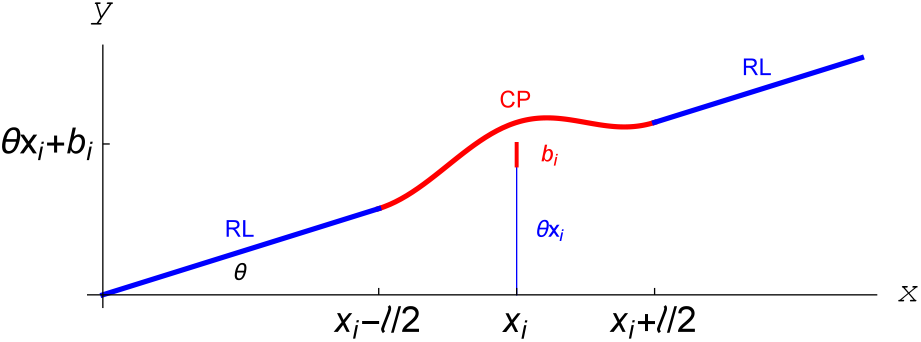
Shape of a cuticular plate when it forms a bulge. It extends from *x_i_* − *ℓ*/1 to *x_i_* + *ℓ*/2 and its average height is *h_i_* = *θx_i_* + *b_i_*. The scales along the *x*- and the *y*-axis are very different.

### D. Tectorial membrane

The TM is visco-elastic. Its Young modulus is in the order of tens to hundreds of kPa and has different properties according the region above which it is located (inner sulcus, RL, or HC) (34). The mechanical properties of the TM are considered to be essential for the tuning ability of the OoC (17, 26). In order to check this assertion, in this study we eliminate the TM motion and replace it by a rigid boundary, located at the constant position *y*_T_(*x*) = 1.

### E. Outer hair cell bundles

We assume that they exert a force that is a function of the tilt angle, which in turn is a function of *h_i_*. We mimic the measured force (35), which has an unstable central region, by means of the expression

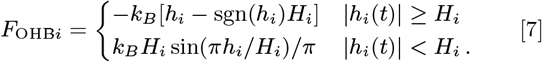

*k_B_* defines the stiffness (we will write *k*_Bolt_ for Boltzmann’s constant) and *H_i_* the range of the unstable region. *F*_OHB*i*_(*h_i_*) is shown in Fig. 4.

**Fig. 4.**
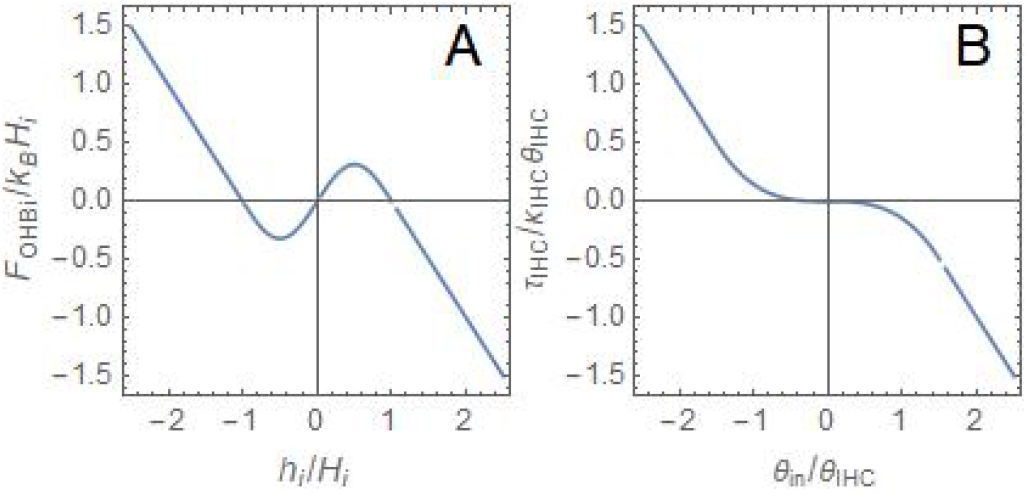
A: Restoring force exerted on the CP by OHB *i*, as a function of the height *h_i_* of the CP over its average position, as stipulated in Section 3E. B: Restoring torque exerted on IHB by the inner hair cell, as a function of the bundle deflection *θ*_in_, as stipulated in Section 3J.

Taking *F*_OHB*i*_ as a function of *h_i_* implies that the work performed by bundle motility vanishes for a complete cycle. Note, however, that if the duration of a cycle is not short compared to the adaptation time (36), *F*_OHB*i*_ becomes historydependent rather than just a function of *h_i_*, and the work that it performs during a cycle does not necessarily vanish.

### F. Outer hair cells

We envision an OHC as a couple of objects, each with mass *m*, connected by a spring. One object is located at the CP and the other at the boundary with the Deiters cell (DC). A special feature of the spring is that its relaxed length can vary. We denote by *c_i_* the contraction of the cell with respect to its resting length, and by *s_i_* the height of the lower object with respect to its average position. We assume that the tension of the OHC has the form

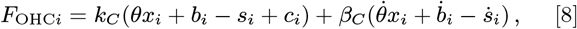

with *k_c_* and *β_c_* positive constant parameters.

The value of *c_i_* is controlled by the inclination of the hair cell bundle. Guided by (28), we assume that when a CP moves towards the TM the hair bundle bends in the excitatory (clockwise) direction. We assume that *h_i_*, scaled with the length *H_i_*, acts as a “degree of excitation,” so that *c_i_* increases with *h_i_*/*H_i_*. Since there must be a maximum length, Δ, by which an OHC can contract, contraction is expected to saturate for large deviations of a CP from its average position. We take this saturation into account by writing

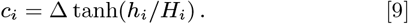

The degree of excitation *h_i_/H_i_* may be identified with *Z*(*X* − *X*_0_)/2*k*_Bolt_*T* in Eq. 3 of (37).

Since *c_i_* is not a function of the distance between the objects on which *F*_OHC*i*_ acts, *F*_OHC*i*_ *can* perform non vanishing work in a complete cycle, as will be spelled out in Section 4C.

### G. Deiters cells

We model a DC as a massless spring that connects the lower object in the OHC to the BM (the mass of the DC is already lumped into *m*). We also include dynamic friction between adjacent lower objects, that encourages oscillation in phase. Denoting by *y*_BM_ the height of the BM above its average position, we write

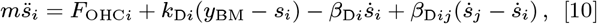

where DC *j* is adjacent to DC *i*. Since DCs are longer for larger *x, k*_D*i*_ and *β*_D*i*_ could depend on *i*.

### H. Hensen cells

We model the HC as a strip with parabolic shape of evenly distributed mass *m*_H_, with its left extreme tangent to the RL and the other extreme pinned at (*x,y*) = (*L* + *L*_HC_, 0). These requirements impose 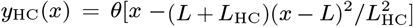. The torque exerted on the HC with respect to the pinning point is 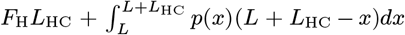, and equals the time derivative of the HC angular momentum, 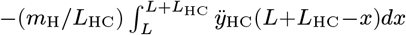, leading to

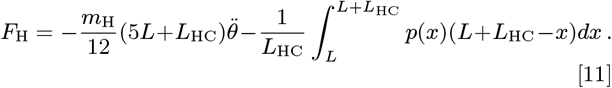

Since we assume that the pressure vanishes in the SM, we replace the upper limit in the integral with the end of the subtectorial channel. We will take this end over the position where the HC has maximum amplitude, namely, 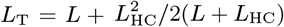.

### I. Inner sulcus

We take the pressure *p*_in_ in the inner sulcus (IS) as uniform and proportional to the increase of area (volume per thickness) with respect to the relaxed IS. We write

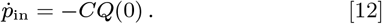

*C* is some average value of the Young modulus divided by the area (in the *xy*-plane) of the soft tissue that coats the IS and *Q*(0) is the flow rate for *x* = 0.

### J. Inner bundle

We locate it at *x* = 0 and assume that its length is almost 1. Models for the torque exerted by the fluid on the IHB abound (10, 27, 38, 39). We will take a simpler approach. The force exerted by viscosity on a segment of the IHB between *y* and *y* + *dy* is proportional to the relative velocity of endolymph with respect to the segment, and we denote it by 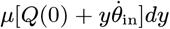, where *μ* is a drag coefficient and we have replaced *υ*(*y*) by its average. On average, the force per unit length is 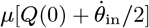. We identify this force with the pressure difference and write

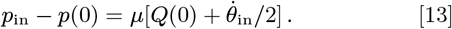

*p*(0) is the pressure at *x* = 0.

The torque exerted by viscosity is 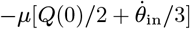. We assume that the moment of inertia of the bundle is negligible and write 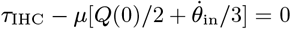, with *τ*_IHC_ the torque exerted by the cell. We assume that the inner hair cell does not rotate, and *τ*_IHC_ is a function of *θ*_in_. It seems reasonable to assume that the IHB does not have a central range with negative stiffness as the OHB, since it could cause sticking of the bundle at any of the angles at which stiffness changes sign. We will assume that, as a remnant of the OHB negative stiffness, *∂τ*_IHC_/*∂θ*_in_ vanishes at *θ*_in_ = 0 [alike Fig. 1(C) in (36)], and write

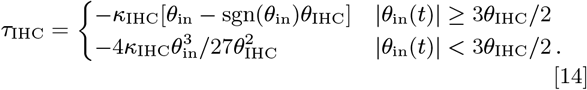

*τ*_IHC_ is a smooth function of *θ*_in_ and the parameters *κ*_IHC_ and *θ*_IHC_ determine its size and the extension of the low stiffness region. *τ*_IHC_(*θ*_in_) is shown in Fig. 4.

We assume that the rate of impulses passed to the auditory nerve is an increasing function of the amplitude |*θ*_in_|.

### K. Basilar membrane

We assume that the BM drives the lower ends of the DCs, all of them by the same amount. In the absence of noise, we take *y*_BM_ = *A* cos*ω*_BM_*t*.

### L. Noise

We investigate the ability of the OoC to filter noise present in the input *y*_BM_; we do not consider noise that arises at the OoC itself. We mimic white noise by adding to *y*_BM_ in Eq. [10] four sinusoidal additions *A*_N_ cos(*ω_j_t* − Φ_*j*_), where the frequencies *ω_j_* are randomly taken from a uniform distribution in the range 0 ≤ *ω_j_* ≤ 2*ω*_BM_. *ω*_1_ (respectively *ω*_2_, *ω*_3_, *ω*_4_) is re-randomized at periods of time 0.7 (repectively 0.9, 1.1, 1.3). The values of Φ*j* are initially random, and afterwards are taken so that *A*_N_ cos(*ω_j_t* − Φ_*j*_) is continuous. is taken so that the average energy added to the DC (for a slice of thickness *D*_0_) is of the order of *k*_Bolt_*T* ~ 4.2 × 10^−21^ J. The initial values of most variables are taken from normal distributions appropriate for average energies of the order of 0.5*k*_Bolt_*T* per degree of freedom; we assume that these initial values become unimportant after a short time.

### M. Procedure

Equations [2] and [4] can be integrated analytically over *x* and, likewise, the integrals of *p* in Eqs. [5], [6] and [11] are performed. After this, using the constitutive relations [7], [9] and [14], we are left with a system of ordinary differential equations for functions of time, that is solved numerically (29).

### N. Parameters

Clearly, parameters vary among species, among individuals, and along the cochlea. We tried to set parameters of reasonable orders of magnitude. The values we took are based on the literature (6, 11, 12, 22, 40–42), when available. When forced to guess, our main guideline was to choose values that lead to large flow for a given amplitude of the input. Additional criteria were fast stabilisation, similar amplitudes of *b*_1_(*t*) and *b*_2_(*t*), avoidance of beating, resonance frequency in a reasonable range, etc. Some of the parameters have almost no influence.

Since bending of the CPs has not been considered in the literature, the value of *k*_CP_ deserves explicit discussion. Since the thickness of the CP’s lip is roughly a third of its length (33), we expect *k*_CP_ to be of the order of the lip’s Young modulus divided by 3^3^. A range of reasonable values for the RL’s Young modulus is stated in (42). *β*-actin and spectrin are relatively scarce in the lip region (33), possibly indicating scarce cross-linking and therefore less resistance to binding; accordingly, we took the Young modulus 50 kPa, close to the lower bound quoted in (42), leading to *k*_CP_ ~ 2 kPa. For *ρ* = 10^3^kgm^−3^, *ν* = 7 × 10^−7^m^2^/s and *D*_0_ = 5 × 10^−6^m, this can be written as 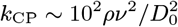.

The parameters we took are listed in Table 1.

**Table 1.** Parameters used in our calculations. We assume that the maximal contraction of the OHC takes its bifurcation value, which for these parameters is Δ_*c*_ = 0.254. The third row indicates the section where the symbol is defined. The system of units is defined in Section 2B.

| Parameter | $L$ | $L_H$ | $x_1$ | $x_2$ | $\ell$ | $m$ | $m_H$ | $I_{RL}$ | $\kappa_{RL}$ | $k_{CP}$ | $\beta_{CP}$ | $k_C$ | $\beta_C$ | $k_{D1}$ | $k_{D2}$ | $\beta_{D1}$ | $\beta_{D2}$ | $\beta_{D12}$ | $k_B$ | $H_1$ | $H_2$ | $\kappa_{IHC}$ | $\theta_{IHC}$ | $C$ | $\mu$ |
| --- | --- | --- | --- | --- | --- | --- | --- | --- | --- | --- | --- | --- | --- | --- | --- | --- | --- | --- | --- | --- | --- | --- | --- | --- | --- |
| Value | 10 | 10 | 3 | 7 | 2 | 10 | 120 | $2 \times 10^3$ | $10^3$ | $10^2$ | 3 | 50 | 43 | 400 | 400 | 3 | 3 | 3 | 10 | $6.5 \times 10^{-3}$ | $5 \times 10^{-3}$ | 10 | $5 \times 10^{-3}$ | 2 | 10 |
| Definition | 2B | 2B | 3C | 3C | 3C | 3C | 3H | 3B | 3B | 3C | 3C | 3F | 3F | 3G | 3G | 3G | 3G | 3G | 3E | 3E | 3E | 3J | 3J | 3I | 3J |

## 4. Results

### A. Main Results

We regard the maximal contraction of the OHC, Δ, as a control parameter, i.e., the parameter that quantifies the power generated within the system.We find that there is a critical value of the control parameter, Δ = Δ_*c*_, such that for Δ > Δ_*c*_ the OoC undergoes self-oscillations (non zero output for zero input), whereas for Δ < Δ_*c*_ it does not. If we take Δ = Δ_*c*_, the OoC becomes a critical oscillator (43–45). Expressions for the output amplitude close to Δ = Δ_*c*_ are worked out in Appendix 3. For the parameters in Table 1, we found Δ_*c*_ = 0.254*D*_0_ and in the limit Δ → Δ_*c*_ the oscillation frequency is *ω_c_* = 5.338 in units of 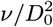. Since these values depend on the parameters we took, they are valid only for the particular considered slice. The length of an OHC is typically ~ 10*D*_0_, so that Δ_*c*_ corresponds to a contraction of a few percent.

Critical oscillator behaviour can be a great advantage for the purpose of tuning and amplitude compression (44). Here we explore the possibility of having this behaviour in the “second filter.” Hence, in the following we study the case Δ = Δ_*c*_. If the input from the BM at the slice position selects a frequency near *ω_c_*, then the OoC would provide additional tuning; if it doesn’t, it would provide an alternative mechanism for tuning. If Δ is moderately close to Δ_*c*_, then the second filter could provide moderate additional tuning and compression.

Figure 5 shows the gain |*θ*_in_|/|*y*_BM_ | as a function of the frequency, for several amplitudes of *y*_BM_. Our results show remarkable similarity between the passage from the BM to the IHB and the experimentally known gain of the BM with respect to the stapes (2, 46). In both cases, weaker inputs acquire larger amplification and tighter selectivity. Except for the case of the lowest amplitude, the gain becomes independent of the amplitude far from the resonance frequency. The inset in Fig. 5 is an expansion of the range 5.1 ≤ *ω*_BM_ ≤ 5.5. It shows that the gains for moderate amplitudes behave as expected from a critical oscillator in the vicinity of the bifurcation point (see Appendix 3). As it usually occurs in critical phenomena, there is also a remarkable similarity between Fig. 5 and Fig. 5b of (23), despite the deep differences between the considered models.

**Fig. 5.**
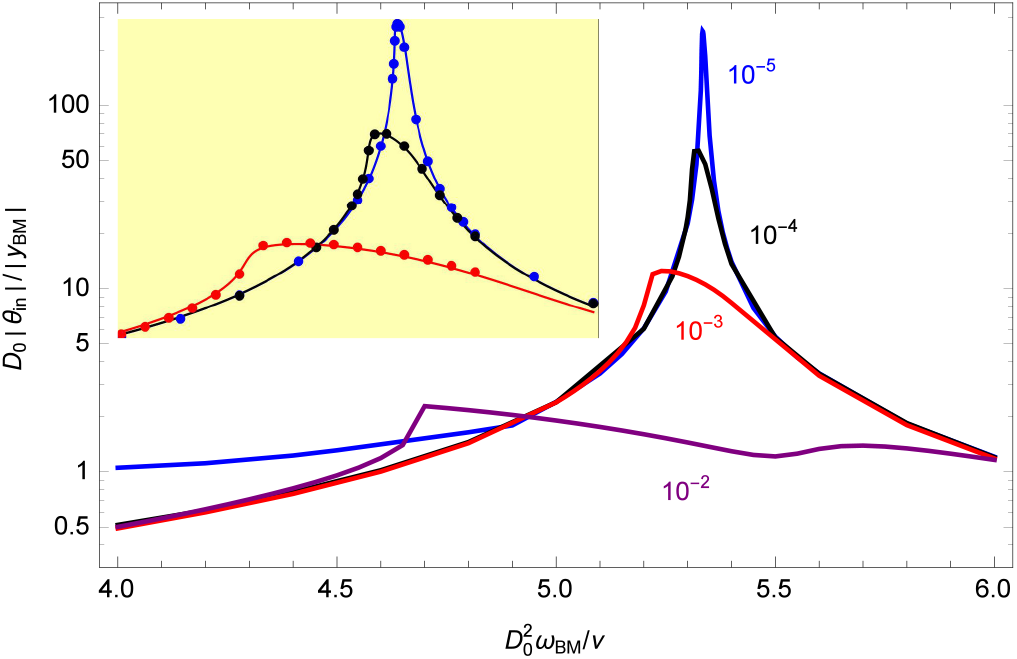
Gain supplied by the OoC. |*θ*_in_| is the root mean square (rms) amplitude of the deflection angle of the IHB and |*y*_BM_| is the rms amplitude of the height of the BM at the point where it touches the DC, 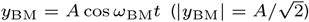. The value of *A* is marked next to each curve. In these evaluations we have ignored thermal noise. Inset: the dots are calculated values for our system and the lines obey Eq. [39] with the fitted values |*B*| = 1.8 × 10^3^, *a* = 6.6 × 10^−4^, *χ*_1_ = −0.66 (for the three lines). Our units are specified in section 2B.

The gain curves are skewed, providing a faster cut at lower frequencies than at higher frequencies. This feature is complementary to the selectivity provided by the cochlear partition, that provides a fast cutoff for high frequencies.

Indeed, early experiments did find that, as the frequency is lowered below resonance, the pressure levels required to excite the auditory nerve or to generate a given electrical response at an inner hair cell grow faster than the pressure levels required to bring about a given vibration amplitude at the BM (47, 48). The credibility of these experiments was limited by the suspicion that the mass or the damage caused by the Mössbauer source or by the reflecting bead used in the measurement of BM vibration could affect its tuning, and also by the large variability (49), which implies that comparison of quantities measured in different individuals may not be justified. A later experiment (18) compared vibrations at a BM site with the response of auditory nerve fibres innervating neighbouring inner hair cells, and obtained good agreement between BM and nerve responses, provided that BM displacements were high-pass filtered, or BM velocities were considered instead. Still, it could be argued that if for faint amplitudes |*θ*_in_| is very sharply tuned, then the spike of the nerve response curve could infiltrate undetected between consecutive measured points. Also, the variability argument could be reversed to claim that the absence of a second filter in a few cases does not rule out its existence in other individuals or locations.

If the transduction from the BM to the IHB has critical oscillator behaviour, then the amplitude compression at resonance of neural activity should be larger than that of BM motion. Indirect experimental support for this scenario is provided by measurements of the OoC potential (50) and of the ratio between the amplitudes of motion of the RL and the BM (51).

Figures 6 and 7 compare the time dependencies of the input and of the output in the case of a small signal when noise is present. The signal had the form *y*_BM_ = *A* cos *ω*_BM_*t* during the periods 2000 < *t* < 4000 and 6000 < *t* < 8000, and was off for 0 < *t* < 2000 and 4000 < *t* < 6000. We took *A* = 3 × 10^−5^ and *ω*_BM_ = 5.329 (which corresponds to the highest gain for this amplitude). Our model for noise is described in Section 3L. The input *y*_total_(*t*) is the sum of the signal and the noise. Panel A in each of these figures shows the entire range 0 < *t* < 8000, and the other panels focus on selected ranges.

**Fig. 6.**
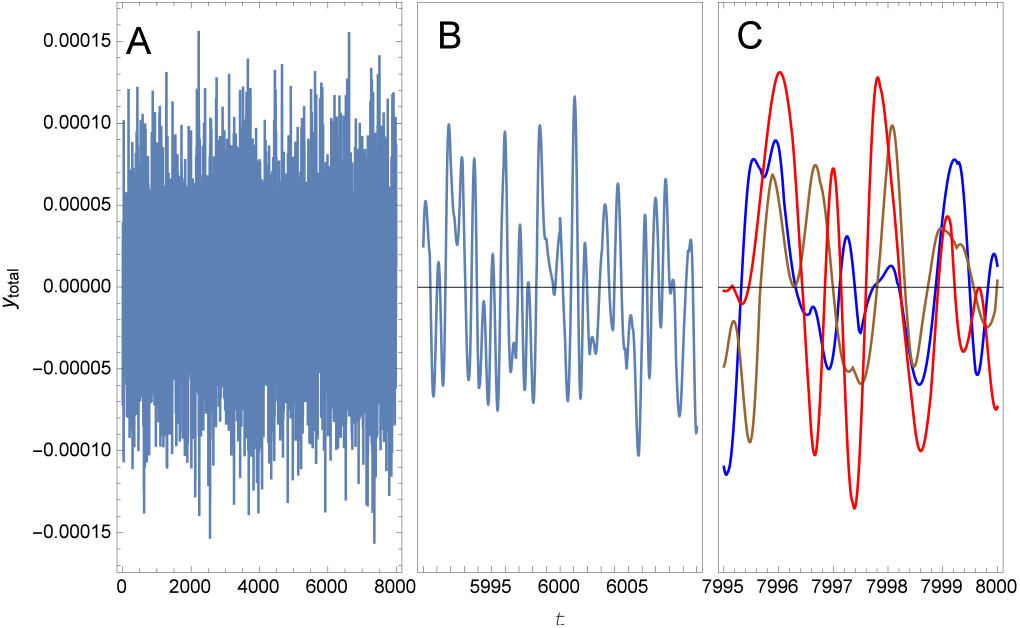
Input when noise is present. The height of the BM relative to its equilibrium position is 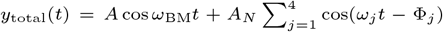, with *A* = 3 × 10^−5^, *ω*_BM_ = 5.329, *A_N_* = 3.5 × 10^−5^, *ω_j_* periodically randomised and Φ_*j*_ determined by continuity. A: Entire considered range. B: Range that contains the instant *t* = 6000, at which the signal is switched on. C: Three lines obtained during equivalent periods while the signal was on: the blue line describes the period 7995 < *t* < 8000 and the brown (respectively red) line describes a lapse of time that preceded by 400 (respectively 3500) times 2*π*/*ω*_BM_. Our units are specified in section 2B.

Figure 6B shows *y*_total_(*t*) in a range such that during the first half only noise is present, whereas during the second half also the signal is on. It is hard to notice that the presence of the signal makes a significant difference. Figure 6C contains three lines: the blue line shows *y*_total_(*t*) during the lapse of time indicated at the abscissa, close to *t* = 8000; the brown line refers to the values of *y*_total_(*t*) at times preceding by 400 × 2*π*/*ω*_BM_ ≈ 472, after the signal had been on during about 1500 time units, and the red line refers to times preceding by 3500 × 2*π*/*ω*_BM_, close to the end of the first stage during which the signal was on. Despite the fact that the signal was identical during the three lapses of time considered, there is no obvious correlation among the three lines.

In contrast to Fig. 6A, we see in Fig. 7A that *θ*_in_ is significantly larger when the signal is on than when it is off. The blue, brown and red lines in Fig. 7B show *θ*_in_(*t*) for the same periods of time that were considered in Fig. 6C. In this case the three lines almost coalesce, and are very close to the values of *θ*_in_(*t*) that are obtained without noise. In particular, we note that the phase of *θ*_in_(*t*) is locked to the phase of the signal.

**Fig. 7.**
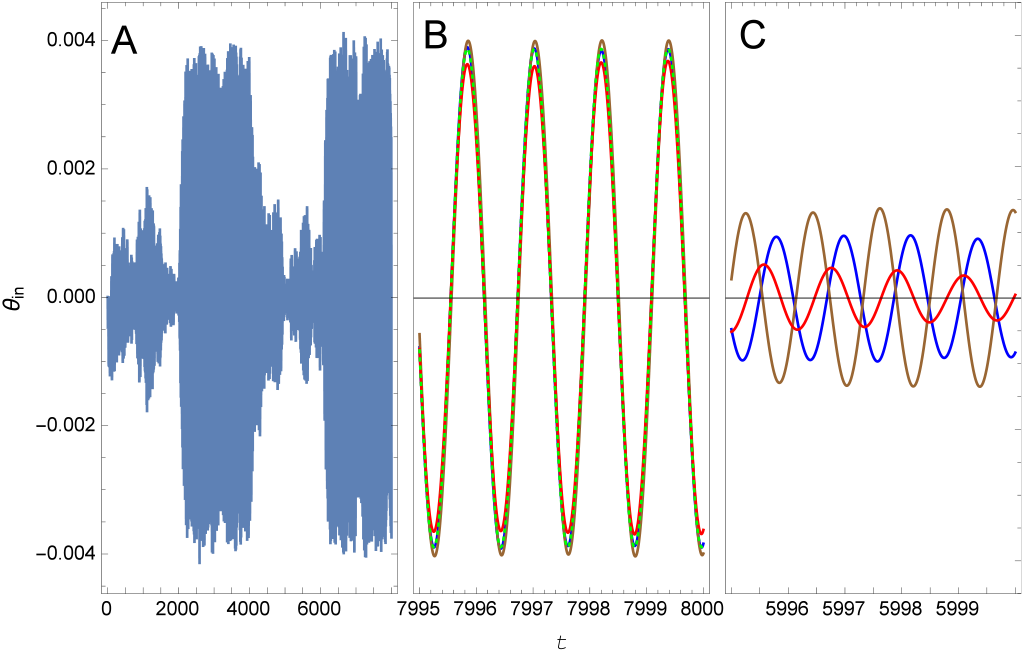
Output, *θ*_in_(*t*), for the situation considered in Fig. 6. A: Entire range. B: The blue, brown and red lines correspond to the same periods of time shown in Fig. 6C; the dotted green line was obtained by dropping the contribution of noise to *y*_total_(*t*). C: The three lapses of time shown in panel B have been shifted 2000 units to the left, so that they cover ranges when no signal was present.

Figure 7C shows *θ*_in_(*t*) for 5995 < *t* < 6000, and also for periods of time preceding by 400 and by 3500 times 2*π*/*ω*_BM_.

In the three cases, the signal was off. We learn from here that the IHB undergoes significant oscillations due to thermal fluctuations even though there is no signal. We also note that there is “ringing,” i.e., oscillations are larger after the signal was on, and it takes some time until they recover the distribution expected from thermal fluctuations. Unlike the case of Fig. 7B, the phase is not locked, and wanders within a relative short time. If the brain is able to monitor the phase of *θ*_in_(*t*), an erratic phase difference between the information coming from each of the ears could be used to discard noise-induced impulses, and a continuous drift in phase difference could be interpreted as motion of the sound source.

Strictly following our models, if the IHB were attached to a fixed point in the TM, it wouldn’t move. In a more realistic model, motion of the BM would tilt the pillar cells, leading to inclination of the IHB. Therefore, in the case of an attached IHB, the signal to noise ratio of the IHB’s inclination would be similar to that of BM motion. On the other hand, comparison of Figs. 6 and 7 shows that the signal to noise ratio of *θ*_in_ is much larger than that of *y*_BM_, strongly suggesting one possible answer to the question of why the IHB is not attached to the TM: in this way the signal to noise ratio increases remarkably.

### B. Motion of each component

Figure 8 shows the amplitudes and phases of *Q*(0)/*ν*, *b*_1,2_, *s*_1,2_ and *Lθ* for a broad range of input frequencies. *b*_1_ and *b*_2_, and likewise *s*_1_ and *s*_2_, nearly coincide, except for a small range of frequencies slightly above the resonance, where motion in the first OHC is considerably smaller than in the second. *L*|*θ*| is roughly three times smaller than |*b*_1,2_| and *θ* is nearly in anti-phase with *b*_1,2_ (lags by ~ 200°). The opposite motions of the RL and the CPs may be attributed to incompressibility and to our assumption of a rigid TM, so that when one of them goes up the other has to go down. *Q*(0) typically lags behind *b*_1,2_ by ~ 80°; following the incompressibility argument, *Q*(0) is positive when the sum of the subtectorial volumes taken by the CPs, the RL and the HC is decreasing. All the variables undergo a 180° change when crossing the resonance.

**Fig. 8.**
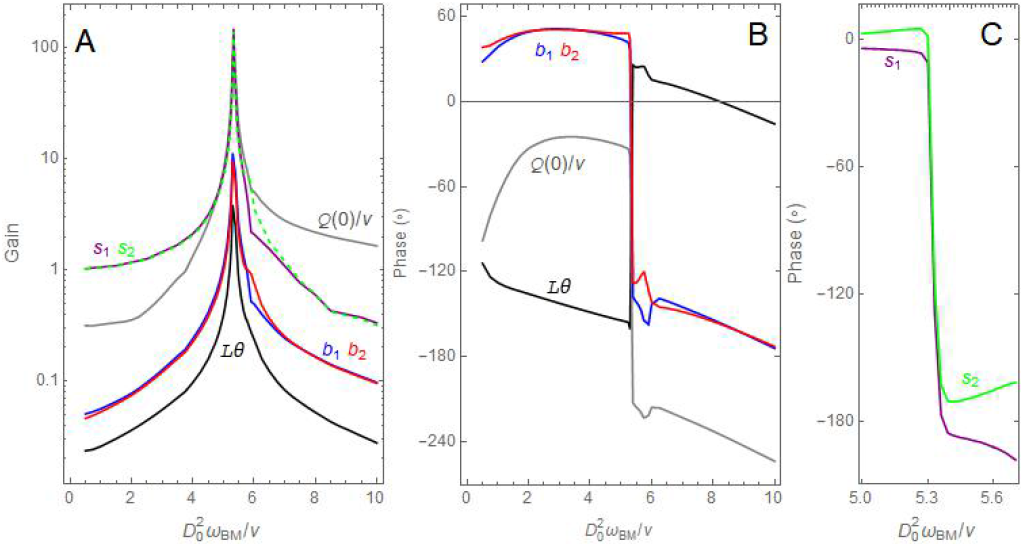
Amplitude and phase of several variables, relative to the input *y*_BM_ = 10^−4^*D*_0_ cos *ω*_BM_*t* (which typically corresponds to ~ 50 dB SPL). A: Amplitude, as defined in Eq. [17]. For visibility, *s*_2_ is depicted by a dashed line. B: Phase by which the variable precedes the input. Phases that differ by an integer number of cycles are taken as equivalent. The phase of a variable is defined as the phase of its first harmonic (see Appendix 1). C: Phases of *s*_1_ and *s*_2_ near the resonance. Here and in the following figures noise has been neglected.

At resonance, |*b*_1>2_| ~ 0.5 × 10^−3^*ℓ*, indicating that the CPs are just moderately bent.

We note that close to the resonance the amplitudes of *s*_1,2_ are larger than those of *b*_1,2_ and *Lθ*. This result is in agreement with the finding of a “hotspot” located around the interface between the OHCs and the DCs, where vibrations are larger than those of the BM or of the RL (52).

Separate motion of the CPs and the RL has not been detected experimentally. We could argue that the lateral spacial resolution of the measuring technique did not distinguish between the CPs and the surrounding RL, so that the measured motion corresponds to some average, but the spot size reported in (24) (less than a *μ*m) excludes this possibility. In the case of (24) there was electrical simulation, and no input from the BM. The most likely possibility is that the TM recedes when the CPs go up, so that the RL does not have to recede and is mainly pulled by the CPs. For a relevant comparison with experiment, the RL motion in Fig. 8 would have to be interpreted as motion relative to the TM, which was within the limits of reproducibility in (24).

A marked difference between (19) and Fig. 8 is the absence of phase inversion when crossing the resonance, possibly indicating that the maximum gain (amplitude of RL motion divided by BM motion) occurs at a frequency beyond the range considered in Fig. 5 of (19) (which includes the maximum of BM motion). A sharp decrease of the phase of the RL relative to the BM occurs in (51).

### C. Mechanical energy transfer

The power delivered by electromotility of OHC *i* is 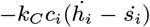. Using Eq. [9] and dropping the terms that give no contribution through a complete cycle, the work performed by electromotility during a complete cycle is

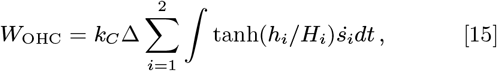

where integration involves a complete cycle. Since both *h_i_* and *S_i_* undergo a phase inversion when crossing the resonance, the sign of *W*_OHC_ remains unchanged.

Similarly, the work per cycle performed by DC *i* on the BM is

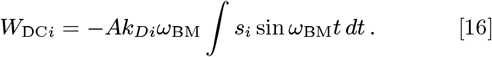

*W*_DC*i*_ > 0 if and only if the phase of *s_i_* is in the range between 0° and 180° (or equivalent). We see from Fig. 8C that very near the resonance *W*_DC1_ and *W*_DC2_ are both negative, indicating that the OoC takes mechanical energy from the BM. Below this region (but still in the range shown in this figure), *W*_DC1_ < 0, *W*_DC2_ > 0, and the opposite situation occurs above this region.

Figure 9 shows the values of these works close to the resonance frequencies, for *A* = 10^−4^ and *A* = 10^−3^. Most of the energy required for motion in the OoC is supplied by electromotility, and a small fraction is taken from the BM.

**Fig. 9.**
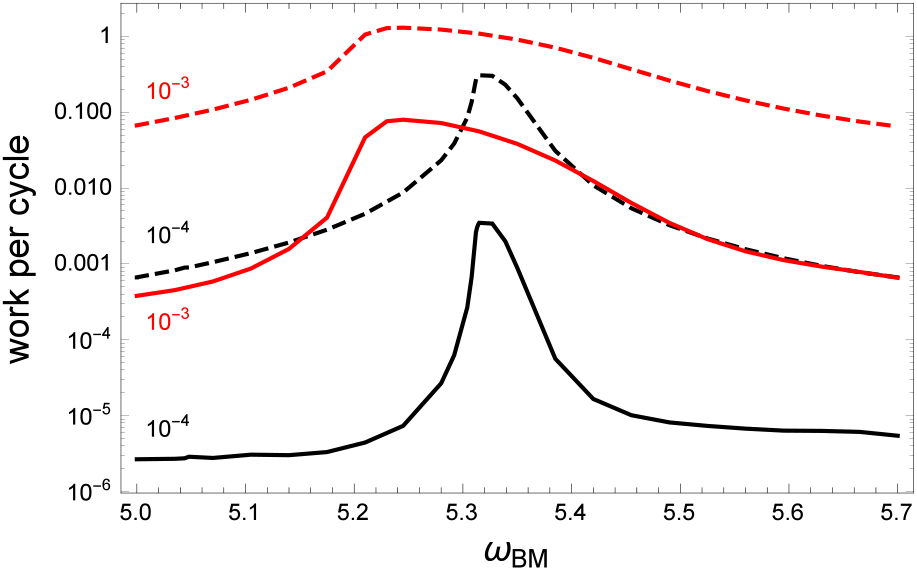
Work performed during a cycle for frequencies close to resonance. The dashed lines refer to the work delivered by electromotility, *W*_OHC_, and the continuous lines to the work taken from the BM, −*W*_DC1_ − *W*_DC2_. *y*_BM_ = *AD*_0_ cos *ω*_BM_*t* and the value of *A* is shown next to each line.

### D. Amplification of the travelling wave

The accepted explanation for active tuning by the cochlea is amplification of each Fourier component of the travelling wave along the segment between the oval window and the place where this component resonates (6, 53), followed by attenuation beyond this place.

In our set of models *W*_DC1_ + *W*_DC2_ is the only exchange of mechanical energy between the considered slice of the OoC and its surroundings; the larger this work, the larger the amplification of the wave. Within a more realistic model, the energy exchange described here should be regarded as a contribution. In the case of Fig. 9, energy is taken from the travelling wave, leading to attenuation.

With the parameters of Table 1, amplification would occur for 5.8 ≲ *ω*_BM_ ≲ 6.4, as shown in Fig. 10. The work performed on the BM depends on the amplitude of *y*_BM_ and can even change sign. If this work is positive/negative the amplitude will increase/decrease, thus approaching the amplitude at which *W*_DC1_ + *W*_DC2_ = 0.

**Fig. 10.**
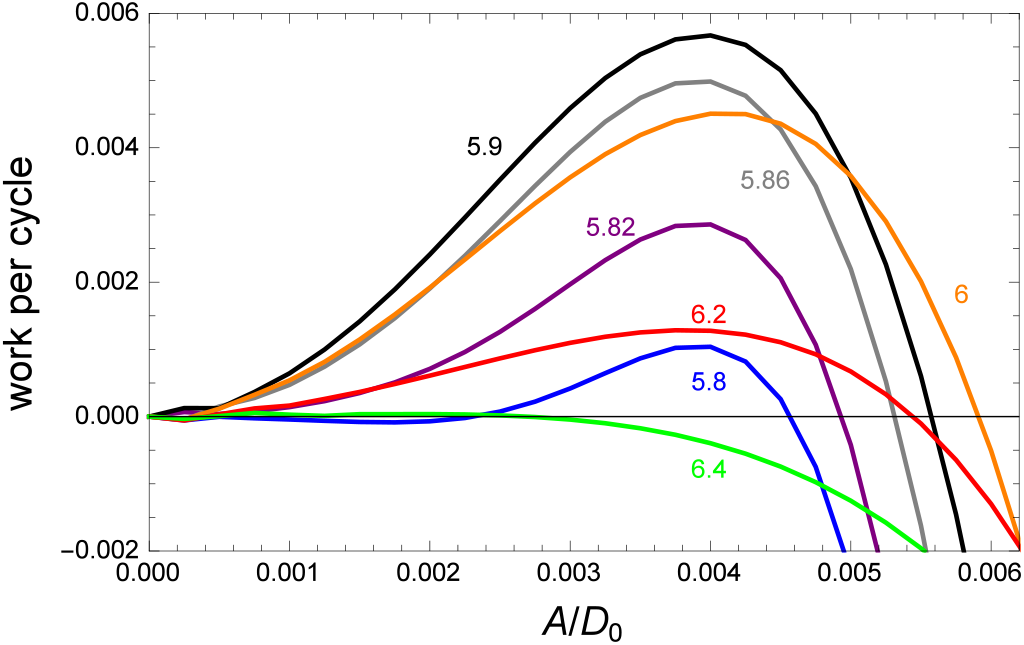
Work performed on the BM as a function of the amplitude of the BM oscillations. The parameter *ω*_BM_ is shown next to each curve. The travelling wave is amplified if this work is positive and attenuated if it is negative. After many cycles, the amplitude of the BM oscillations would be largest for *ω*_BM_ ≈ 6

Contrary to the accepted explanation, the amplification range in Fig. 10 lies above *ω_c_*. This could be the case if the resonance of the “first filter” lies above that of the second, but the situation can also change if the parameters are slightly varied. For example, if we raise *k*_D2_ by 10%, to 440, Δ_*c*_ becomes 0.290, *ω_c_* becomes 5.506, and the travelling wave is amplified in the range 4.9 < *ω*_BM_ < 5.4, as shown in Fig. 11. Conceivably, the advantage of having several OHCs per slice (rather than a single stronger OHC) is the possibility of adjusting the resonance frequencies of both filters, so that they cooperate rather than interfere with each other.

**Fig. 11.**
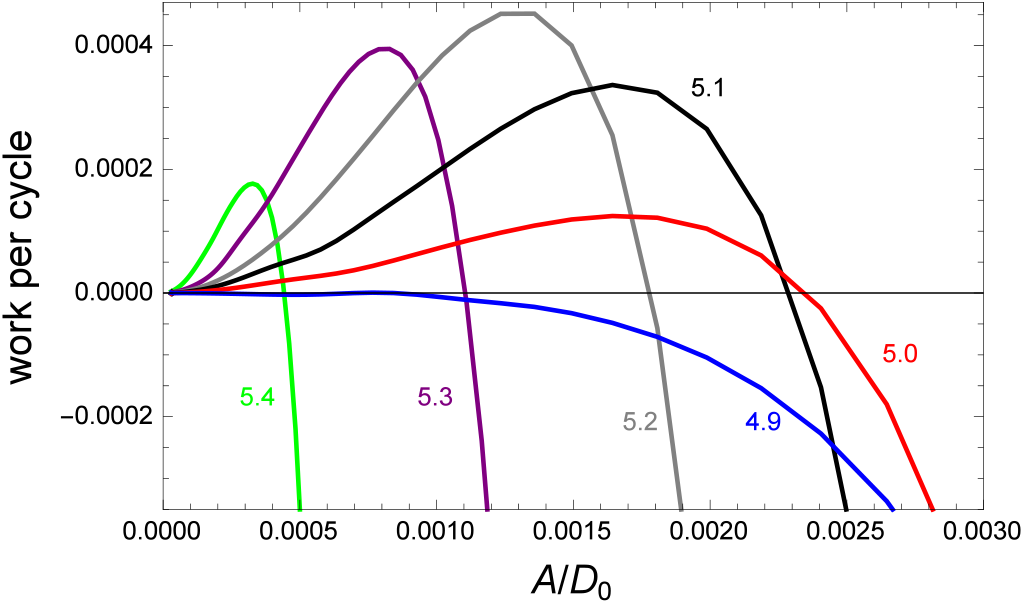
Like Fig. 10, this time for *k*_*D*2_ = 440 rather than 400.

Figures 10 and 11 indicate that the travelling wave is attenuated for frequencies below the considered range. However, we should note that the energy transferred for given work per cycle is not proportional to the travelled distance, but rather to the travelling time. Therefore, the largest influence will be that of the slices where the travelling wave is slow, close to the resonance of the first filter. The number of cycles that the travelling wave is expected to undergo while passing through a given region is estimated in Appendix 4. Dependence of amplification on time rather than on distance could help to understand the unexpected results obtained in (53).

### E. Time dependence of the output

Figure 12 shows *θ*_in_(*t*) for *A* = 10^−5^ and frequencies near resonance. The blue envelope was obtained at resonance frequency, *ω_R_* = 5.334, the pink envelope at *ω*_BM_ = 5.34 and the green envelope at *ω*_BM_ = 5.32. In the case of resonance, the output amplitude raises monotonically until a terminal value is attained. Out of resonance, the amplitude starts increasing at the same pace as at resonance, overshoots its final value, and then oscillates until the final regime is established.

**Fig. 12.**
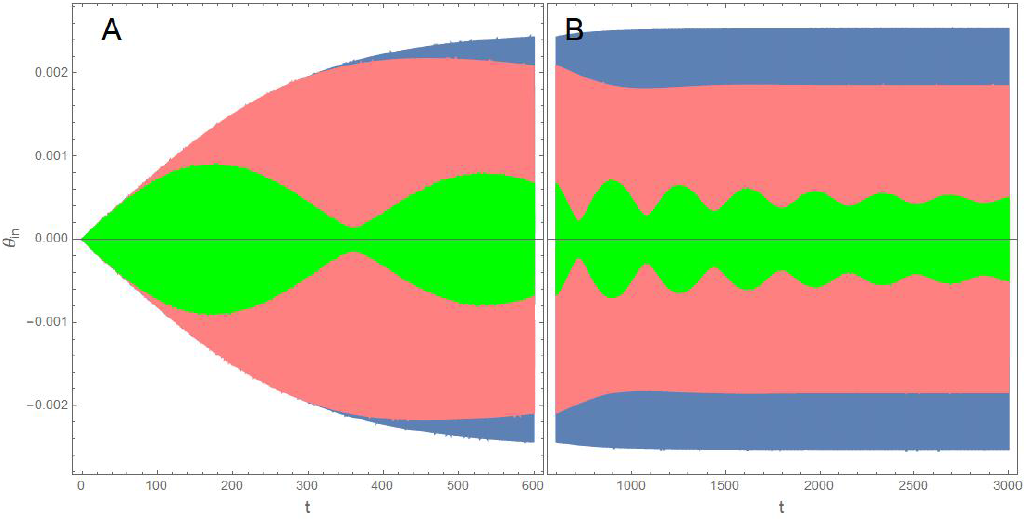
Angle of the IHB as a function of time in response to *y*_BM_ = 10^−5^*D*_0_ cos*ω*_BM_*t*. Blue: resonance frequency, *ω*_BM_ = *ω_R_* = 5.334; pink: *ω*_BM_ = 5.34; green: *ω*_BM_ = 5.32. A: 0 ≤ *t* ≤ 600. B: *t* ≥ 600. *ω_R_* approaches the critical frequency *ω_c_* in the limit of small amplitude.

This initial behaviour implies that the IHB will start reacting to the input if *ω*_BM_ is moderately close to *ω_R_*, before it can tell the difference between these two frequencies. Conversely, for a given *ω*_BM_, there will be several slices of the OoC with a range of frequencies *ω_R_* close to *ω*_BM_ that will start reacting to this input. As an effect, all these slices will send a fast alarm telling that something is happening, before it is possible to discern the precise input frequency.

In contrast with a forced damped harmonic oscillator, when out of resonance, motion of the OoC does not assume the frequency of the input even after a long time, but is rather the superposition of two modes, one with the input frequency *ω*_BM_, and the other with the resonance frequency *ω_R_*. If *ω*_BM_ = (*n*_1_/*n*_2_)*ω_R_*, where *n*_1,2_ are mutually prime integers, then the motion has period 2*n*_2_*π*/*ω_R_*. Figure 13 shows *θ*_in_(*t*) for *ω*_BM_ = (2/3)*ω_R_* and for *ω*_BM_ = (4/3)*ω_R_*.

**Fig. 13.**
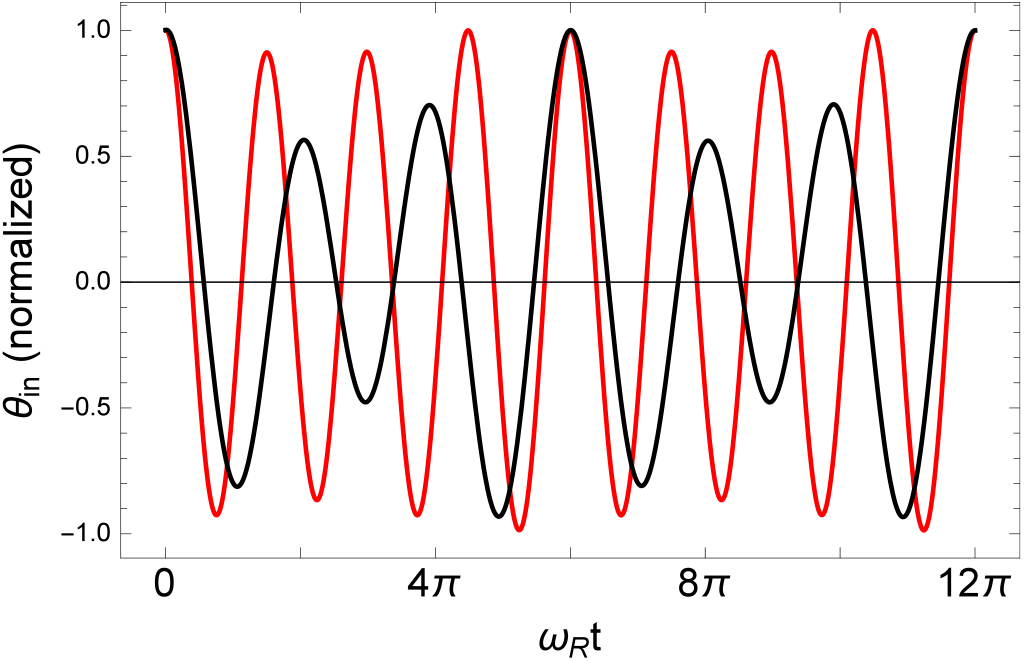
*θ*_in_(*t*) during a short period of time. Black: *ω*_BM_ = (2/3)*ω_R_* red: *ω*_BM_ = (4/3)*ω_R_. t* is the time elapsed after a maximum of *θ*_in_, roughly 4000 time units after the input was set on. *A* = 10^−5^.

### F. Nonlinearity

We studied the deviation from sinusoidality of *θ*_in_(*t*) at resonance frequency, when the periodic regime is established. Writing 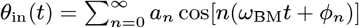, the even harmonics vanish. Taking the origin of time such that *φ*_1_ = 0, we found the values in Table 2.

**Table 2.**
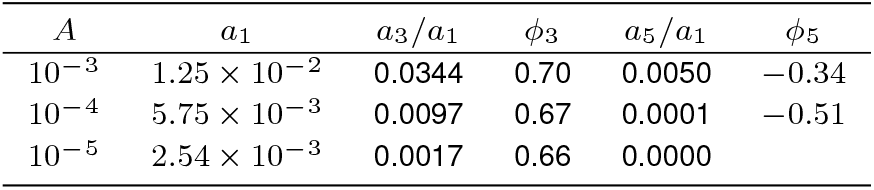
*θ*_in_ ≈ *a*_1_ cos *ω*_BM_*t* + *a*_3_ cos[3(*ω*_BM_*t* + *φ*_3_) + *a*_5_ cos[5(*ω*_BM_*t* + *φ*_5_)] *A* is the peak value of the input and *ω*_BM_ equals the resonance frequency. *φ*_3,5_ are the phases with respect to the first harmonic of *θ*_in_.

## 5. Discussion

We have built a flexible framework that enables to test many possibilities for the mechanical behaviour of the components of the OoC. The models we used imply that even by taking the basilar membrane motion as an input, the OoC can behave as a critical oscillator, thus providing a second filter that could enhance frequency selectivity and improve the signal to noise ratio. This framework can be used to explore and theoretically predict different effects that would be hard to observe experimentally. Although the models considered here are oversimplifications, they enabled us to obtain features that are compellingly akin to those observed in the real OoC.

According to our models, the reason for fluid flow at the IHB region is the vertical motion of the CPs, the RL and the HC, but other drives are also possible (54, 55). Flow could be due to shear between the TM and the RL, squeezing of the IS, or deviation of part of the RL from the *x*-axis, implying an *x*-component of its velocity when it rotates.

For comparison of the relative importance of each of these mechanisms, we examine the peak values that we obtained for *A* = 10^−4^ at resonance frequency. For *Q*(0), which in our units equals the average over *y* of *υ*(*y*), we found ~ 2 × 10^−2^. The vertical velocity of the CPs is less than 10^−2^. From here we expect that the shear velocity of the RL with respect to the TM will be less than that, and the average fluid velocity even smaller.

The peak value of 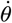 is ~ 5 × 10^−4^. Assuming that the length of the RL that invades the IS is ~ 4*D*_0_, squeezing would cause a flux rate of ~ 10^−3^. It therefore seems that the mechanism that we have considered is the most important, providing a sort of self-consistency check. In the case of a flexible TM, *θ* would be larger and the flux due to squeezing would grow accordingly.

The following sections describe examples of possible modifications of our models. Some of them we have already studied and others have not been studied thus far.

### A. Bundle motility

Bundle motility can be eliminated from the model by setting *H_i_* = 0 in Eq. [7] (but not in [9]). We still obtain that the OoC can behave as a critical oscillator, but the critical value for OHC contraction rises to Δ_*c*_ = 0.262. Our conclusion is thus that bundle motility helps attainment of critical oscillator behaviour, but is not essential.

### B. Removal of the HC

This was done by setting *L*_T_ = *L* and *F*_H_ = 0. The bifurcation value of Δ increased to Δ_*c*_ = 0.273, suggesting that an advantage of the HC is reduction of the amount of contraction required to achieve criticality. The comparison may be somewhat biased by the fact that our parameters were optimised with the HC included.

### C. Natural extensions

In order to describe a situation as it occurs in nature, our models should consider flexibility of the TM. A model with TM that just recedes would be easy to implement, but a realistic model that includes shearing should also allow for motion of the base of the IHB.

Our models could deal with the longitudinal dimension along the cochlea (*z*) by taking an array of slices, with parameters and input *y*_BM_ that are functions of *z*. The interaction between neighbouring slices could be mechanical, mediated by the phalangeal processes and deformation of the TM, or hydrodynamic, mediated by flow along the IS and the SM.

For simplicity, in Eq. [9] *c_i_* is an odd function of *h_i_*. In reality, OHCs contract by a greater amount when depolarised than what they elongate when hyperpolarised. [9] corresponds to the assumption that there are equal probabilities for open and for closed channels (37). We have found that the asymmetry between contraction and elongation is essential for demodulation of the envelope of a signal, as it occurs in (50).

Instead of adding elements to the set of models, an interesting question is how much can be taken away and still have a critical oscillator. We can show that a system of two particles, with a “spring” force between them of the form [8] that depends on the position of one of the particles, and with appropriate restoring and damping coefficients, behaves as a critical oscillator with an unusual bifurcation diagram. The critical control parameter of this “bare” oscillator (with the same parameters used in Table 1) is considerably smaller than the value of Δ_*c*_ that we found for the OoC. These bare oscillators (one for each OHC) drive the entire OoC.

## ACKNOWLEDGMENTS

We are indebted to Anders Fridberger, David Furness, Karl Grosh, James Hudspeth, Daibhid Maoiléidigh, Yehoash Raphael and Luis Robles for their answers to our inquiries.

## Appx1: Periodic non sinusoidal functions

### A. Amplitude

The amplitude of a periodic, or approximately periodic, function *f* will be defined as the root mean square deviation from its average,

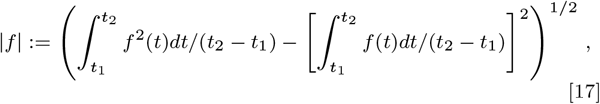

where *t*_2_ − *t*_1_ is an integer number of periods.

### B. Phase differences

We consider two real functions, *f*_1_(*t*) and *f*_2_(*t*), that have the same period 2*π/ω*. We define the ‘phase’ *φ* of *f*_2_ with respect to *f*_1_ by the value that maximises the overlap between these functions when the time is advanced in *f*_1_ by *φ/ω*, i.e., by the value that maximises ∮ *f*_1_(*t* + *φ/ω*)*f*_2_(*t*)*dt*.

Equivalently, if we write 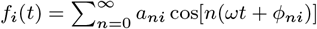, we have to maximize 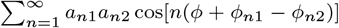, implying 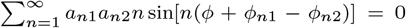. We note that a dc component in any of the functions has no influence on the phase. If *f*_1_(*t*) and *f*_2_(*t*) have the same shape, then *φ*_*n*1_ − *φ*_*n*2_ is independent of *n* and *φ* = *φ*_12_ − *φ*_11_.

In the case of quasi-sinusoidal functions, such that |*a*_*n*1_*a*_*n*2_/*a*_11_*a*_12_| < *ϵ* ≪ 1 for *n* > 1, we look for a solution *φ* = *φ*_12_ − *φ*_11_+*O*(*ϵ*). We expand sin[*n*(*φ*+*φ*_*n*1_ − *φ*_*n*2_)] = sin[*n*(*φ*_12_ − *φ*_11_ + *φ*_*n*1_ − *φ*_*n*2_)] + *n* cos[*n*(*φ*_12_ − *φ*_11_ + *φ*_*n*1_ − *φ*_*n*2_)](*φ* − *φ*_12_ + *φ*_11_) + *O*(*ϵ*^2^) and obtain

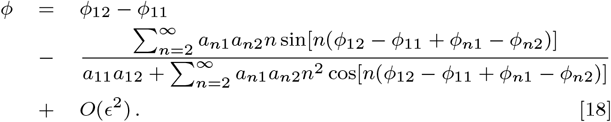

In this article *f*_1_(*t*) is proportional to cos*ωt*, so that the phase depends solely on the first harmonic of *f*_2_(*t*) and becomes

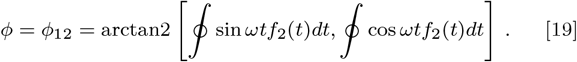

We note that the phase is not additive, i.e., the phase of *f*_3_ with respect to *f*_1_ not necessarily equals the phase of *f*_2_ with respect to *f*_1_ plus the phase of *f*_3_ with respect to *f*_2_.

## Appx2: Fluid flow in a narrow channel with small rapid wall motion

The channel is defined by *T* = {(*x,y*)| 0 < *x* < *L*, *ξ*(*x,t*) < *y* < *D*_0_}. The flow problem is characterised by three non-dimensional parameters:

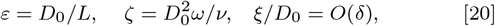

where 2*π/ω* is the oscillation period (in time) of *ξ*, and *ν* ~ 1 mm^2^/s is the kinematic viscosity. Typical values for the length parameters above are

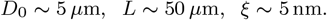

Thus, *ε* ~ 0.1, while *δ* ~ 10^−3^. We shall work under the canonical scaling *ζ* = *αε*, where *α* = *O*(1).

The fluid velocity (*υ, u*) and pressure *p* satisfy the time-dependent Stokes equation:

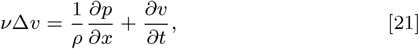

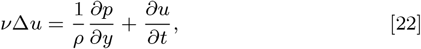

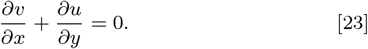

Here Δ is the Laplacian operator. No-slip boundary conditions are assumed on the channel’s lateral boundary.

To convert the problem to a nondimensional formulation we scale (*υ, u*) by 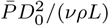, where 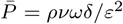 is the scale for *p*. We further scale *x* by *L, y* by *D*_0_, and time by 1/*ω*. Finally, we introduce the scaling *ξ_t_* = *δD*_0_*ωη_t_*, where *η*(*x,t*) is dimensionless and the subscript denotes derivative. Substituting all of this into the fluid equations, and retaining the original notation for the scaled variables, we obtain

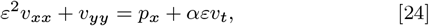

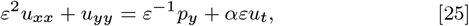

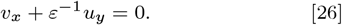

### First order expansion

We expand *υ* = *υ*^0^ + *ευ*^1^ + … and similarly for *p, u*, and the flux 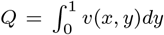. To leading order *p*^0^ = *p*^0^(*x, t*), and *u*^0^ = *u*^0^(*x, t*) due to [25] and [26]. However, the no-slip boundary conditions imply *u*^0^ = 0. To leading order in *δ* the horizontal motion of the wall is negligible up to *ε*^3^, and we retain only the vertical motion. Therefore, the kinematic boundary condition at *y* = 0 is

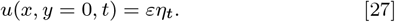

The leading order term *υ*^0^ satisfies 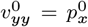 with boundary conditions *υ*^0^(*x*, 0, *t*) = *υ*^0^(*x*, 1, *t*) = 0. Therefore,

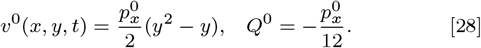

Integrating the incompressibility equation [26] over (0,1), and since to leading order *u* = *εu*^1^, we obtain

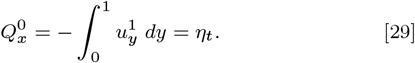

Combining equations [28] and [29] provides an equation for the pressure 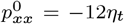. Given the boundary motion *η*(*x,t*), this equation, together with boundary conditions for *p*^0^, can be solved to find the pressure and from it the velocity *υ*^0^ and the flux *Q*^0^.

### Second order expansion

Since *u*^0^ = 0, it follows from equation [25] that also *p*^1^ satisfies *p*^1^ = *p*^1^ (*x, t*). At the next order we obtain

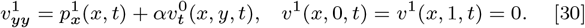

Using equation [28], *υ*^0^ can be expressed in the alternative form *υ*^0^(*x,y,t*) = −6*Q*^0^(*x,t*)(*y*^2^ − *y*). Solving equation [30] for *υ*^1^ we find

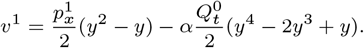

Integrating *υ*^1^ over (0, 1) we obtain

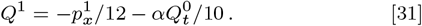

Addition of [28] and [31] gives the following equation, exact up to *O*(*ε*):

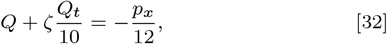

which is equivalent to equation [4].

Similarly, up to *O*(*ε*), *υ*(*x, y, t*) = −6*Q*(*x, t*)(*y*^2^ − *y*) − *ζQ_t_*(*x, t*)(5*y*^4^ − 10*y*^3^ + 6*y*^2^ − *y*)/10. We recall that *Q*(*x, t*) is available from the solution of the system of differential equations in our code. Once *υ*(*x, y, t*) is known, *u* can be obtained from [23] and the boundary conditions, and the full equations [21] and [22] can be checked for self consistency. We have found that while the expansion above was carried out for values of *ζ* smaller than 1, numerical evidence indicates that equation [32] is valid for much larger values of *ζ*. For instance, we consider a representative problem with *ζ* ~ 5. Expansion up to *O*(*ε*) entirely drops *υ_xx_* when evaluating *p_x_* in [24]. Support for this approximation can be based on Fig. 14, where we see that *υ_xx_* is significantly smaller than *p_x_*. Similarly, Fig. 15 shows that *p* is essentially independent of *y*.

**Fig. 14.**
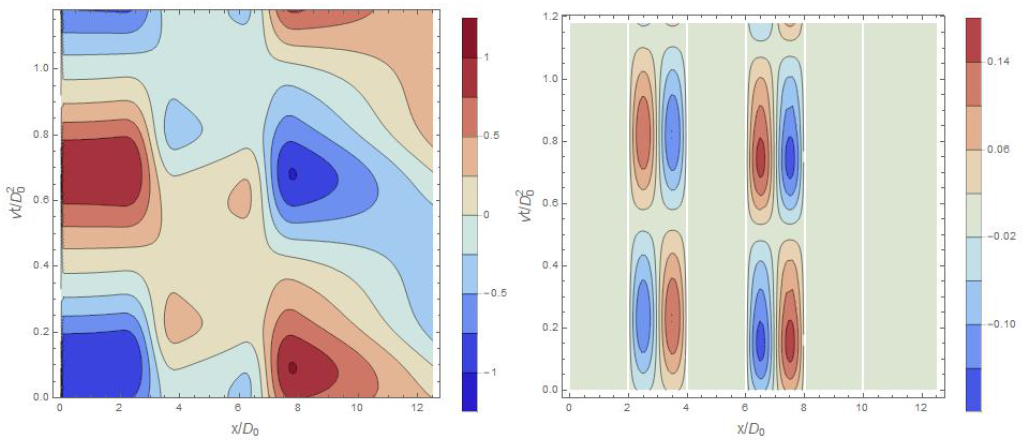
Left: Contour plot of normalized pressure gradient, 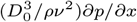, as a function of position and time, obtained using [32] and thus neglecting *ε*^2^*υ_xx_* in [24]. Right: *y*-average of the neglected term, 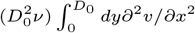. The white lines are places where *υ_xx_* is discontinuous. The time span describes one cycle, beginning and ending when *θ* assumes its most negative value. For disambiguation, all quantities in the legends and in this caption are dimensional. The color scale bars are different for each graph.

**Fig. 15.**
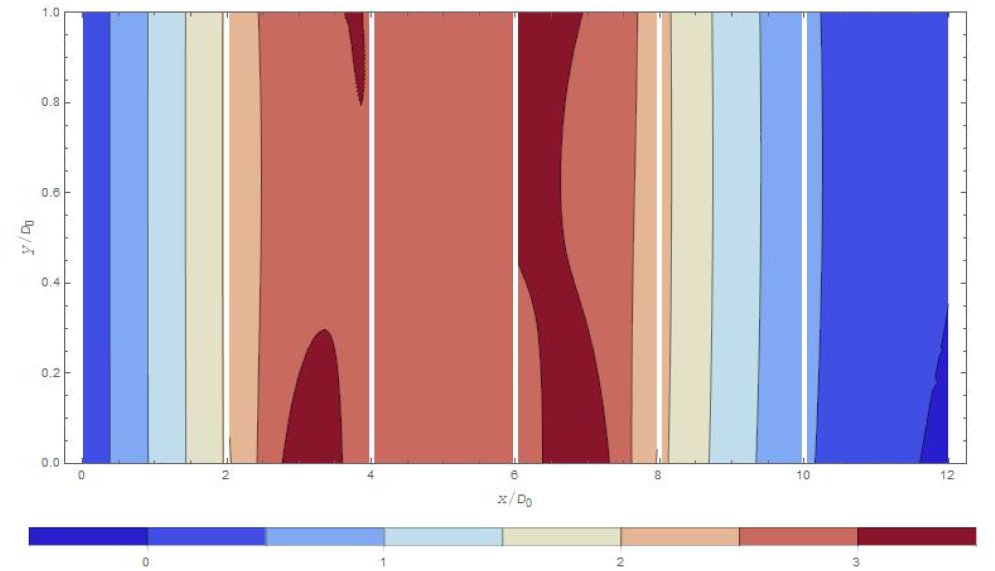
Pressure *p*(*x, y, t* = 1.2*π/ω*_BM_) in the in the subtectorial channel. The pressure unit in the color scale bar is 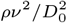. At the moment depicted in this snapshot the RL is moving downwards and the CPs are moving upwards. At the white lines the pressure is discontinuous, but since the *y*-dependence is small the discontinuity is not visible in the figure. For *t* ≠ 1.2*π/ω*_BM_, |*p*(*x, y* = 0.5*D*_0_, *t*) − *p*(*x, y* = 0, *t*)| is typically smaller.

## Appx3: Critical Oscillators

Let us deal with an oscillator in which the signal *Y* can be expressed in terms of the response *X* in the form

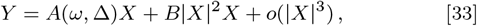

such that *A*(*ω_c_*, Δ_*c*_) = 0. (*ω_c_*, Δ_*c*_) is called “bifurcation point.” Let us write Ω = *ω* − *ω_c_, δ* = Δ − Δ_*c*_ and assume that *B* can be approximated as constant and *A* can be expanded as

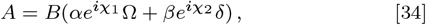

with *α, β* > 0 and 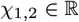.

In order to have a spontaneous response without any signal, *αe*^*iχ*1^ Ω + *βe*^*iχ*2^ *δ* + |*X*|^2^ has to vanish. In this case, from the imaginary part we obtain

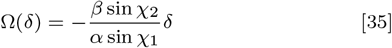

and then, from the real part,

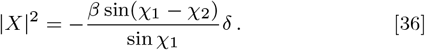

Equation [36] indicates that non-vanishing spontaneous responses occur either for *δ* > 0 or for *δ* < 0, depending on whether the signs of sin(*χ*_1_ − *χ*_2_) and sin *χ*_1_ are opposite or the same. In our case, Δ is the maximal contraction of the OHCs and spontaneous responses were found for *δ* > 0.

Let us now consider forced oscillations, *Y* ≠ 0. From [33] and [34] we have

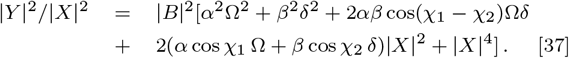

In particular, for Δ = Δ_*c*_,

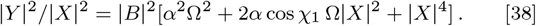

In our case the signal is the deviation of the BM from its equilibrium position, the response is the inclination of the IHB, and [38] predicts the gain

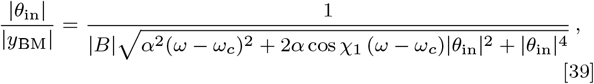

where *B, α* and *χ*_1_ do not depend on *ω* or |*y*_BM_|.

For small amplitudes and close to the bifurcation point, and for appropriately fitted values of Δ_*c*_, *ω_c_*, |*B*|, *α, β, χ*_1_ and *χ*_2_, our results are in good agreement with Eqs. [35], [36] and [39].

## Appx4: Number of cycles during which the travelling wave is amplified/attenuated

We want to estimate the number of cycles *n_cy_* experienced by a wave of frequency *ω*BM as it travels across the region *z*_1_ ≤ *z* ≤ *z*_0_, where *z*_0_ is the position (distance from the oval window) of the slice we consider and *z*_1_ is the position where the wave starts to be amplified or attenuated significantly.

The dispersion relation can be obtained from Eqs. (2.17) and (2.40) (neglects damping) in (6):

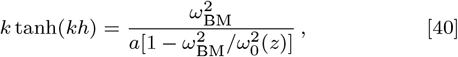

where *k* is the wave number, *h* the height of the chamber above or below the partition, *a* is a constant and *ω*_0_(*z*) is the first-filter resonant frequency at position *z*.

For *kh* ≪ 1 and *ω*_BM_ ≪ *ω*_0_(*z*), [40] becomes 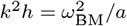, and therefore *a* = *V*^2^(0)/*h*, where *V*(0) is the speed of the travelling wave in the long wavelength limit. For *ω*_BM_ close to *ω*_0_(*z*), *kh* is significantly larger than 1 and [40] becomes

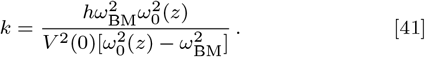

The number of cycles is 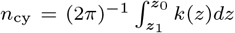. Assuming that *dw*_0_/*dz* = −λ*w*_0_ with constant λ, and using [41] we obtain

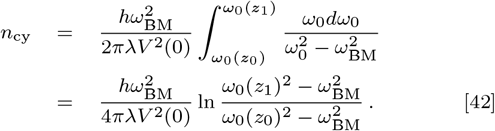

Taking *h* = 0.0005m, *ω*_BM_ = 2*π* × 5kHz, λ = 150m^−1^ (56) and *V*(0) = 15m/s (57), we obtain 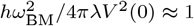.

## References

1. Dallos P (1992) The Active Cochlea. J Neurosci 12 4575–4585

2. Robles L, Ruggero MA (2001) Mechanics of the Mammalian Cochlea. Physiol Rev 81 1305–1352

3. Hudspeth AJ (2008) Making an Effort to Listen: Mechanical Amplification in the Ear. Neuron 59 530–545

4. Puria S, C R Steele CR (2008) Mechano-Acoustical Transformations. The senses: A comprehensive reference, eds Basbaum AI et al (Academic, NY) Vol 3, pp 166–201

5. Hudspeth AJ (2014) Integrating the active process of hair cells with cochlear function. Nat Rev Neurosci 15 600–614

6. Reichenbach T, Hudspeth AJ (2014) The physics of hearing: fluid mechanics and the active process of the inner ear. Rep Prog Phys 77 076601

7. Manley GA, Gummer AW, Popper AN, Fay RR, eds (2017) Understanding the Cochlea (Springer, Cham)

8. von Békésy G (1960) Experiments in Hearing (McGraw Hill,NY)

9. Peterson LC, Bogert BP (1950) A Dynamical Theory of the Cochlea. J Acoust Soc Am 22 369—381

10. Steele CR, Puria S (2005) Force on inner hair cell cilia. Int J Solids Struc 42 5887–5904

11. Ramamoorthy S, Deo NV, Grosh K (2007) A mechano-electro-acoustical model for the cochlea: Response to acoustic stimuli. J Acoust Soc Am 5 2758–2773

12. Ni G, Elliott SJ, Baumgart J (2016) Finite-element model of the active organ of Corti. J R Soc Interface 13: 20150913

13. Liu Y, Gracewski SM, Nam J-H (2017) Two passive mechanical conditions modulate power generation by the outer hair cells. PLoS Comput Biol 13 e1005701

14. Sasmal A, Grosh K (2019) Unified cochlear model for low- and high-frequency mammalian hearing. Proc Natl Acad Sci USA 116 13983–13988

15. Evans EF, Wilson JP (1975) Cochlear tuning properties: concurrent basilar membrane and single nerve fiber measurements. Science 190 1218–1221

16. Allen JB, Neely ST (1992) Micromechanical models of the cochlea. Physics Today 45 40–47

17. Gummer AW, Hemmert W, Zenner (1996) Resonant tectorial membrane motion in the inner ear: Its crucial role in frequency tuning. Proc Natl Acad Sci USA 93 8727–8732

18. Narayan SS, Temchin AN, Recio A, Ruggero MA (1998) Frequency Tuning of Basilar Membrane and Auditory Nerve Fibers in the Same Cochleae. Science 282 1882–1884

19. Chen F, Zha D, Fridberger A, Zheng J, Choudhury N, Jaques SL, Wang RK, Shi X, Nutall AL (2011) A differentially amplified motion in the ear for near-threshold sound detection. Nat Neurosci. 14 770–774

20. Lee HY, Raphael PD, Xia A, Kim J, Grillet N, Applegate BE, Ellerbee Bowden AK, Oghalai JS (2016) Two-Dimensional Cochlear Micromechanics Measured In Vivo Demonstrate Radial Tuning within the Mouse Organ of Corti. J Neurosci 36 8160–8173

21. Dallos P (2003) Organ of Corti Kinematics. J Assoc Res Oto 4 416–421

22. Nam J-H, Fettiplace R (2010) Force Transmission in the Organ of Corti Micromachine. Biophys J 98 2813–2821

23. Maoiléidigh DÓ, Jülicher F (2010) The interplay between active hair bundle motility and electromotility in the cochlea. J Acoust Soc Am 128 1175–1190

24. Nowotny M, Gummer AW (2011) Vibration responses of the organ of Corti and the tectorial membrane to electrical stimulation. J Acoust Soc Am 130 3852–3872

25. Richter C-P, Quesnel A (2006) Stiffness properties of the reticular lamina and the tectorial membrane as measured in the gerbil cochlea, Auditory Mechanisms: Processes and Models, eds Nutall AL, Ren T, Gillespie P, Grosh K, de Boer E (World Scientific, Singapore) pp 70–78

26. Zwislocki JJ, Kletsky EJ (1979) Tectorial membrane: a possible effect on frequency analysis in the cochlea. Science 204 639–641

27. Nowotny M, Gummer AW (2006) Nanomechanics of the subtectorial space caused by electromechanics of cochlear outer hair cells. Proc Natl Acad Sci USA 103 2120–2125

28. Fridberger A, Tomo I, Ulfendahl M, Boutet de Monvel J (2006) Imaging hair cell transduction at the speed of sound: Dynamic behavior of mammalian stereocilia. Proc Natl Acad Sci USA 103 1918–1923

29. https://www.notebookarchive.org/models-for-organ-of-corti--2020-01-2aqmevw/

30. Tilney LG, Derosier DJ, Mulroy MJ (1980) The organization of actin filaments in the stereocilia of cochlear hair cells. J Cell Biol 86 244–259

31. Raphael Y, Athey BD, Wang Y, Lee MK, Altschuler RA (1994) F-actin, tubulin and spectrin in the organ of Corti: comparative distribution in different cell types and mammalian species. Hear Res 76 173–187

32. Kachar B, Battaglia A, Fex J (1997) Compartmentalized vesicular traffic around the hair cell cuticular plate. Hear Res 107 102–112

33. Furness DN, Mahendrasingam S, Ohashi M, Fettiplace R, Hackney CM (2008) The dimensions and composition of stereociliary rootlets in mammalian cochlear hair cells: comparison between high- and low-frequency cells and evidence for a connection to the lateral membrane. J Neuroscience 28 6342–6353

34. Gueta R, Barlam D, Shneck RZ, Rousso I (2006) Measurement of the mechanical properties of isolated tectorial membrane using atomic force microscopy Proc Natl Acad Sci USA 103 14790–14795

35. Martin P, Mehta AD and Hudspeth AJ (2000) Negative hair-bundle stiffness betrays a mechanism for mechanical amplification by the hair cell. Proc Natl Acad Sci USA 97 12026–31

36. Tinevez J-Y, Jülicher F, Martin P (2007) Unifying the Various Incarnations of Active Hair-Bundle Motility by the Vertebrate Hair Cell. Biophys J 93 4053–4067

37. Fettiplace R, Kim KK (2014) The physiology of mechanoelectrical transduction channels in hearing. Physiol Rev 94 951–986

38. Freeman DM, Weiss TF (1988) The role of fluid inertia in mechanical stimulation of hair cells, Hear Res 35 201–208

39. Sasmal A, Grosh K (2018) The Competition between the Noise and Shear Motion Sensitivity of Cochlear Inner Hair Cell Stereocilia. Biophys J 114 474–483

40. Evans BN, Dallos P (1993) Stereocilia displacement induced somatic motility of cochlear outer hair cells. Proc Natl Acad Sci USA 90 8347–8351

41. Ramamoorthy S, Nuttall AL (2012) Outer hair cell somatic electromotility in vivo and power transfer to the organ of Corti. Biophys J 102 388–398

42. Zagadou BF, Barbone PE, Mountain DC (2014) Elastic properties of organ of Corti tissues from point-stiffness measurement and inverse analysis. J Biomech 47 1270–1277

43. Duke T, Jülicher F (2008) Critical Oscillators as Active Elements in Hearing. Active Processes and Otoacoustic Emissions, Springer Handbook of Auditory Research, eds Manley GA, Popper AN, Fay RR (Springer, New York), pp 63–92

44. Hudspeth AJ, Jülicher F, Martin P (2010) A critique of the critical cochlea: Hopf—a bifurcation—is better than none. J Neurophysiol 104 1219–1229

45. Ó Maoiléidigh D (2018) Multiple mechanisms for stochastic resonance are inherent to sinusoidally driven noisy Hopf oscillators. Phys Rev E 97 022226

46. Johnstone BM, Patuzzi R, Yates GK (1986) Basilar membrane measurements and the travelling wave. Hearing Research 22 147–153

47. Sellick PM, Patuzzi R, Johnstone BM (1983) Comparison between the tuning properties of inner hair cells and basilar membrane motion. Hearing Research 10 93–100

48. Ruggero MA, Rich NC, Recio A, Narayan SS, Robles S (1997) Basilar-membrane responses to tones at the base of the chinchilla cochlea. J Acoust Soc Am 101 2151–2163

49. Rhode WS (1978) Some observations on cochlear mechanics. J Acoust Soc Am 64 I58–176

50. Nuttall AL, Ricci AJ, Burwood G, Harte JM, Stenfelt S, Cayé-Thomasen P, Ren T, Ramamoorthy S, Zhang Y, Wilson T, Lunner T, Moore BCJ, Fridberger A, (2018) A mechanoelectrical mechanism for detection of sound envelopes in the hearing organ. Nature Commun 9: 4175

51. Ren T, He W, Kemp D (2016) Reticular lamina and basilar membrane vibrations in living mouse cochleae. Proc Natl Acad Sci USA 113 9910–9915

52. Cooper NP, Vavakou A, van der Heijden M (2018) Vibration hotspots reveal longitudinal funneling of sound-evoked motion in the mammalian cochlea. Nature Com 9 3054

53. de Boer E, Nuttall AL, Hu N, Zou Y, Zheng J (2005) The Allen-Fahey experiment extended. J Acoust Soc Am 117 1260–1266

54. Nowotny M, Gummer AW (2006) Nanomechanics of the subtectorial space caused by electromechanics of cochlear outer hair cells. Proc Natl Acad Sci USA 103 2120–2125

55. Guinan JJ Jr (2012) How are inner hair cells stimulated? Evidence for multiple mechanical drives. Hearing Research 292 35–50

56. Greenwood DD (1990) A cochlear frequency-position function for several species-29 years later. J Acoust Soc Amer 87 2592–2605

57. Lighthill J (1981) Energy flow in the cochlea. J Fluid Mech 106 149–213

